# Implication of polymerase recycling for nascent transcript quantification by live cell imaging

**DOI:** 10.1101/2023.09.25.559364

**Authors:** Olivia Kindongo, Guillaume Lieb, Benjamin Skaggs, Yves Dusserre, Vincent Vincenzetti, Serge Pelet

## Abstract

Transcription enables the production of RNA from a DNA template. Due to the highly dynamic nature of transcription, live-cell imaging methods play a crucial role in measuring the kinetics of this process. For instance, transcriptional bursts have been visualized using fluorescent phage-coat proteins that associate tightly with mRNA stem loops formed on nascent transcripts. To convert the signal emanating from a transcription site into meaningful estimates of transcription dynamics, the influence of various parameters on the measured signal must be evaluated. Here, the effect of gene length on the intensity of the transcription site focus was analyzed. Intuitively, a longer gene can support a larger number of transcribing polymerases, thus leading to an increase in the measured signal. However, measurements of transcription induced by hyper-osmotic stress responsive promoters display independence from gene length. A mathematical model of the stress-induced transcription process suggests that the formation of gene loops that favor the recycling of polymerase from the terminator to the promoter can explain the observed behavior. One experimentally validated prediction from this model is that the amount of mRNA produced from a short gene should be higher than for a long one as the density of active polymerase on the short gene will be increased by polymerase recycling. Our data suggest that this recycling contributes significantly to the expression output from a gene and that polymerase recycling is modulated by the promoter identity and the cellular state.

**Take away:**

- Quantification of stress-induced promoter transcription dynamics using a live assays reporter system displays no dependence of signal intensity with gene length.
- Mathematical modeling predicts that the formation of gene loops leading to the recycling of polymerases can explain the observed behavior.
- More prevalent polymerase recycling on short genes results in a higher transcriptional output.

## Introduction

mRNA molecules are essential intermediates between acting cellular proteins and information encoded in DNA. The production of mRNA is an intricate process that involves the interaction between numerous complexes to initiate transcription, elongate the transcript and finally produce a matured mRNA molecule ^1,2^. Biochemical analyses have allowed characterization of the various complexes implicated in the production of mRNA ^3–5^. Advances in sequencing have highlighted the diversity of generated mRNA ^6^. At the single cell level, fluorescence in-situ hybridization (FISH) with fluorescent oligonucleotide probes has illustrated the great variability in mRNA content existing between individual cells and suggested the presence of bursts in transcriptional activity^7^. More recently, the development of single cell transcriptomics in yeast^8–10^ has enabled the analysis of the whole transcriptome from individual cells as well as the visualization of the heterogeneity in the transcription profile of each cell.

Although very informative, the drawback of these techniques is that they provide only snapshot characterization of the highly dynamic mRNA production process. To complement these datasets, two different approaches have been devised to image mRNA in living cells. First, fluorescently tagged phage-coat proteins that recognize specific mRNA stem loops were engineered (Figure 1A)^11,12^. Second, RNA aptamers that become florescent when blinding to a specific chemical compound were designed ^13–15^.

**Figure 1.**
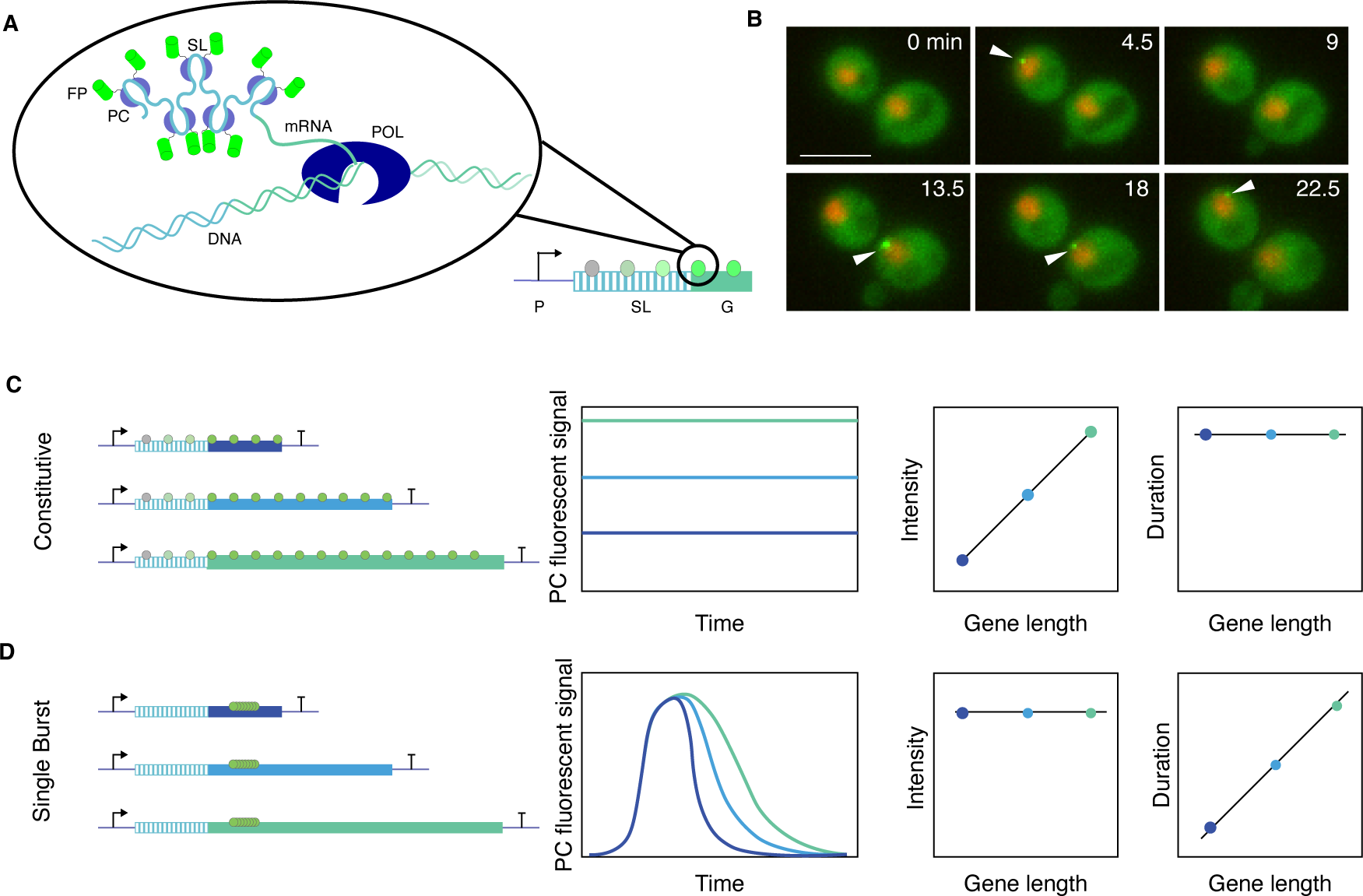
Scheme representing the quantification of nascent mRNA using phage-coat proteins. A. The stem loops (SL) transcribed by the polymerase (POL) and bound by two fluorescently tagged (FP) phage-coat proteins (PC). In panels C and D, the polymerase loaded by the promoter (P) and transcribing the stem loop and the gene body (G) is depicted by a circle. The color of the circle represents the amount of PC bound to the nascent mRNA. B. Images of PP7-GFP marking the nascent transcripts generated by a constitutively active promoter (p*GPD1*). The nuclei are labeled in red using Hta2-mCherry. The arrow heads indicate the presence of a transcription site. The scale bar represents 5µm. C. and D. Expected influence of gene length on the quantification of the transcription site intensity signal monitored with phage-coat proteins for an ideal constitutive promoter (C) or a promoter generating a single burst of transcription (D). In the first case, a dependence of the intensity of the transcription site with gene length is expected, while in the second case, it is the duration of the signal that will increase with gene length.

Live-cell imaging of mRNA has been used for numerous purposes. Initially, this strategy allowed monitoring individual mRNA trafficking between cellular sub-compartments ^11^. It has since been used to directly quantify the production of mRNA by quantifying changes in fluorescence intensity at the transcription site (Figure 1B) and thereby extract key parameters of the transcriptional dynamics ^16^. Numerous properties of the transcriptional process have been characterized thanks to this approach. Importantly, bursts of gene expression have been directly visualized and quantified^16–19^. The interplay between transcription factor binding events and transcription initiation has been observed^20^. Splicing dynamics have been monitored^21^. Additionally, the various parameters extracted from these measurements have been used to build mathematical models of transcription^22,23^.

As these reporter systems become more widely used, it becomes increasingly important to understand how gene architecture controls the intensity and localization of fluorescently labeled mRNA. Obviously, major efforts have been invested in revealing the influence of promoters on the dynamics and level of transcription^24,25^. Sequences controlling the localization and trafficking of the mRNA molecules have been identified^11,26^ and export from the nucleus to the cytoplasm has been monitored^27,28^. However, it has also been noted that the addition of the stem loops and the phage coat protein binding can perturb the trafficking or half-life of mRNAs^29,30^. In this work, we will focus on the effect that gene length can have on the measurement of nascent transcription and total transcriptional output.

To study the transcription induced by a promoter of interest, the promoting sequence is placed upstream of stem loop motifs followed by an open reading frame (ORF) region that can span various lengths and completed by a terminator sequence (Figure 1C). At an ideal constitutive promoter, the loading and transcription of the locus by individual polymerases happen at a constant rate. When the mRNA stem loops are followed by a short ORF, only a few polymerases will be transcribing the locus simultaneously. Thus, the measured transcription site intensity will be lower than on a long gene placed under the control of the same promoter, where a larger number of polymerases can accumulate on the locus and transcribe it simultaneously. Therefore, to a first approximation, a linear dependency between gene length and the intensity of the transcription site is expected (Figure 1C).

Conversely, if we consider a promoter that generates a single burst of transcription, a group of polymerases will be loaded and transcribe the ORF synchronously (Figure 1D). The maximum intensity of the fluorescence signal ought to be independent of gene length. However, the duration of the signal will increase proportionally to gene length, since transcription on the short gene would be completed faster than on the long one.

Native promoters likely fall somewhere between these two extreme cases. Small bursts of transcription will encompass a wide range of dynamics. Therefore, for short genes, one would expect to identify more individual pulses of lower intensity. Instead, on longer genes, these temporally close bursts will be averaged out resulting in a longer and brighter transcription site intensity signal.

To test the relationship between signal intensity and transcript length, we generated transcriptional reporters of various lengths controlled by inducible promoters. Interestingly, some of the reporters tested displayed no dependence between signal intensity and transcript length. Using a simple mathematical model of transcription, we tested the transcriptional parameters that could influence this relationship. These simulations, along with our experimental data, suggest that polymerase recycling enhanced by the formation of gene loops contributes significantly to the transcriptional output by increasing the polymerase load on small genes compared to long ones.

## Materials and Methods

### Strains and plasmids

A list of strains and plasmids used in this study can be found in Tables S1 and S2. All the strains originate from the *Saccharomyces cerevisiae* W303 background^31^. The nuclear marker was generated by fluorescently tagging Hta2 with an mCherry protein^32^. The strain was subsequently transformed with a PP7-GFPenvy expressed from the constitutive ADH promoter^25^. In this strain, the PP7 stem loops controlled by the promoter of interest were integrated in the GLT1 locus^16,25^. The promoters were sub-cloned from previously generated plasmids^25,33^. The fusion between the core of the AGA1 promoter (-150 to 0) and the STL1 upstream activation sequence (-800 to -163) was generated by In-Fusion cloning (Takara Bio) into a plasmid containing the 24xPP7 stem loops. To modulate the transcript length, different homology regions in the GLT1 gene were cloned in the PP7-stem loop containing plasmid (Table S3) to shorten the generated transcript. The insertion of the full 24xPP7 stem loops motif was verified by PCR to eliminate transformants where the stem loop array was increased or diminished because these variations have a strong impact on the intensity of the transcription site.

The measurement of the elongation speed was performed with constructs in which the 24xPP7 stem loops were followed by 24xMS2 stem loops. The 2k or 4k spacers were generated by inserting the sequence of *Schizosaccharomyces pombe* FUS1. The plasmids were transformed in a MAT**α** strain with a HTA2 locus marked with a tdiRFP::NAT cassette and expressing the MS2-GFPenvy construct. The integrities of the PP7 and MS2 loops were verified in this strain that was subsequently mated with a MAT**a** strain producing the PP7-mCherry construct and harboring a HTA2-tdiRFP::TRP. The diploid cells generated by crossing of the MAT**a** and MAT**α** strains and MAT**a/α** cells were selected on SD-T + NAT medium.

The effect of transcript length on the protein level was studied using dynamic protein synthesis translocation reporters (dPSTR)^33^. These constructs consist of two transcription units. First, a constitutively expressed fluorescent protein (R: mCherry, Y: mCitrine) flanked by a small SynZip peptide, which can form strong heterodimers. Second, an inducible peptide that contains a degradation tag, a nuclear localization sequence, and the complementary SynZip peptide^34^. To increase gene length, various portions of the *S. pombe* FUS1 sequence were inserted between the coding sequence of the inducible peptide and the CYC1 terminator. The plasmids p*STL1*-dPSTR-Y with variable lengths were transformed in the same strain bearing a Hta2-CFP nuclear marker and a short p*STL1*-dPSTR-R construct.

### Time-lapse microscopy

The cells were inoculated in SD-full medium (Complete CSM DCS0031, ForMedium) and grown overnight to saturation. The culture was then diluted in fresh medium and maintained in log-phase growth (OD < 0.4) for 24 hours by successive dilutions before imaging. Two hundred microliters of a cell suspension at OD 0.04 were loaded in the well of a 96-well plate (PS96B-G175, SwissCI) coated previously with Concanavalin A (L7647, Sigma-Aldrich). The imaging was performed on a Nikon Ti2 inverted microscope enclosed in a temperature incubation chamber set at 30°C and controlled by micro-manager^35^. The fluorescent excitation light was provided by a Lumencor Spectra III light source.

For the transcription site measurements, the LED intensity was lowered to 5% of the maximum power to minimize photobleaching. Cells were imaged with a 40X oil objective, a quadruple band dichroic (DAPI/FITC/Cy3/Cy5, F68-400, Chroma) and appropriate emission filters. The images were acquired with a Hamamatsu ORCA-Fusion sCOMS camera. Using a piezo stage (Nanodrive, Mad City Lab City), five Z-planes were recorded (-1 to +1 µm) in the fluorescent channels where the transcription sites were recorded every 10 to 20 seconds. The nuclear marker and the bright field image recorded to segment the cells were only imaged every third time point. At every time points, up to four XY-positions per well were visited and the accuracy of the focal plane was ensured using the hardware autofocus. Before the fourth time point, the acquisition was paused to allow the addition of 100µl of stimulation medium concentrated threefold.

For the dPSTR imaging, the LED intensity was lowered to 50% of the maximum. Cells were imaged with a 40X air objective, a quadruple band dichroic (CFP/YFP/RFP/Cy7, F68-037, Chroma) and appropriate emission filters. Five fields of view per well for all illumination channels were recorder every 2 to 5 minutes.

### Flow chamber

Osmotic stress pulses were generated in a flow chamber (μ-slide VI 0.4, Ibidi). Cells were grown, as described previously, to OD 0.2 before loading and were attached to the bottom of the channel using a coating of Concanavalin A. Two media, SD-full and SD-full + 0.4M NaCl + Alexa680-dextran (D34681, Molecular Probes), were output towards the flow chamber via flow generated by pressure controlled reservoirs (LineUp FlowEZ, Fluigent). These media were mixed in calculated ratios to generate specific changes in flow chamber NaCl concentration by a pulse-width modulation scheme with a duty cycle of 500ms using solenoid valves and an Arduino Uno controller^25,36^. Changes in NaCl concentration over time corresponded proportionally to the Alexa680-dextran dye concentration.

### Image analysis

The recorded time-lapse measurements were analyzed using the YeastQuant platform^37^. The bright field segmentation was performed with CellPose based on the cyto2 model^38,39^. These detected cells were combined with an intensity segmentation of the nuclear marker in order to define a nucleus and a cytoplasm object for each cell. Since the transcription site is often found at the periphery of the nucleus, the nucleus object was enlarged by a 5-pixel wide ring to improve detection of transcriptional events. Based on the 40x objective, each pixel represents 0.1625 µm and the objects possess the following radii: Nucleus: 0.7 ± 0.15 µm, ExpandedNucleus: 1.4 ± 0.15 µm, Cell: 2.6 ± 0.5 µm.

The difference between the intensity of the mean of the ten highest pixels in this expanded nucleus object and the median intensity of the object is used as a proxy for the intensity of the transcription site. The maximum intensity of the transcription site is calculated between the average of the maximum of the trace and its two neighboring time points and the mean intensity of the first three time points before the stimulus. The detection of a transcriptionally active cell is based on the connection between these ten high intensity pixels. If at least five of these pixels are locally connected to each other, an active transcription site is detected. In a single cell trace, at least four consecutive active transcription site have to be present to define that a single cell is transcribing. The first and last frames when these active transcription sites are detected are used to define the Start and Stop times of transcription, thereby allowing to define the duration of transcription.

### RT-qPCR

Cells were cultivated in YPD (YEP broth, CCM0405, ForMedium) overnight to saturation. They were subsequently grown for 24hrs in log-phase by multiple dilutions to obtain 30 ml of culture at OD 0.4. For the reference sample, five milliliters of culture were used. Then, the remaining 25 ml were stressed with 1.6 ml of 5M NaCl to reach a final concentration of 0.4M. At specific time points, 5 ml of culture were removed for RNA extraction and the cells were harvested by centrifugation. Following resuspension in 750µl Trizol, 150µl chloroform was added. After homogenization and centrifugation at 4°C, the aqueous phase was recovered. The mRNAs were precipitated with isopropanol and washed with Ethanol 70% before resuspension in water. The reverse transcription was performed to generate the cDNA (SuperScript IV VILO, Invitrogen). The cDNA was diluted ten times and 3µl were used as template to perform the qPCR (LightCycler 480 SYBR Green Master mix and LightCycler 480II, Roche) using the primers listed in Table S4. The constitutive gene VCX1 was used as a reference gene^40^, STL1 and CTT1 were used as control for stress-inducible genes. The primers to monitor the GLT1 abundance were selected in the last 500 bp of the ORF which is present in all the constructions. The mRNA abundance for each time point was estimated by calculating the 2^(Cp_REFGene)^/2^(Cp_TargetGene)^.

### Mathematical Model

The model of transcription was established using the SimBiology toolbox in Matlab (R2021b). The reactions depicted in Figure S3 were modeled by mass action kinetics. The reaction rates and additional parameters of the model are listed in Table S5. Thousand stochastic runs are performed with the model for a total simulated time of 45 min. At time = 5 minutes, the Stress is set to 100. When simulating a pulse, the Stress is set back to zero at time = 10 minutes. A linear interpolation of the stochastic model results is performed to obtain regularly spaced time points every second. The temporal evolution of the PP7 signal is calculated for each time point as the sum of the products between the number of polymerases on each segment of the gene and the value of the fluorescent signal generated by this segment. This fluorescent signal rises from 0 to 1 for the first 1.5 kB of the transcript where the stem loops are encoded and then stays at one for all the other segments. The number of transcripts is equivalent to the number of polymerases that initiate transcription.

## Results

The influence of gene length on the transcriptional process was monitored using modified versions of the 24xPP7-stem loop construct from Larson^16^. The original construct has two homology regions in the GLT1 locus: in the promoter and at the start of the gene, thereby allowing the insertion of an auxotrophy marker, a promoter of interest, and the 24xPP7 stem loops between these two sequences. This locus was selected because GLT1 is one of the longest genes in the genome and is non-essential. In this study, we modulated the length of the transcript by selecting 4 additional homology regions inside the GLT1 ORF to shorten the transcript from 7.9 kB (composed of 1.5 kB for the stem loops and 6.4 kB from the gene) to 2.2 kB (Figure 2A). This strategy enables retention of the same genomic environment for all constructs and direct monitoring of the effects of transcript length on the quantified transcription site fluorescence.

**Figure 2.**
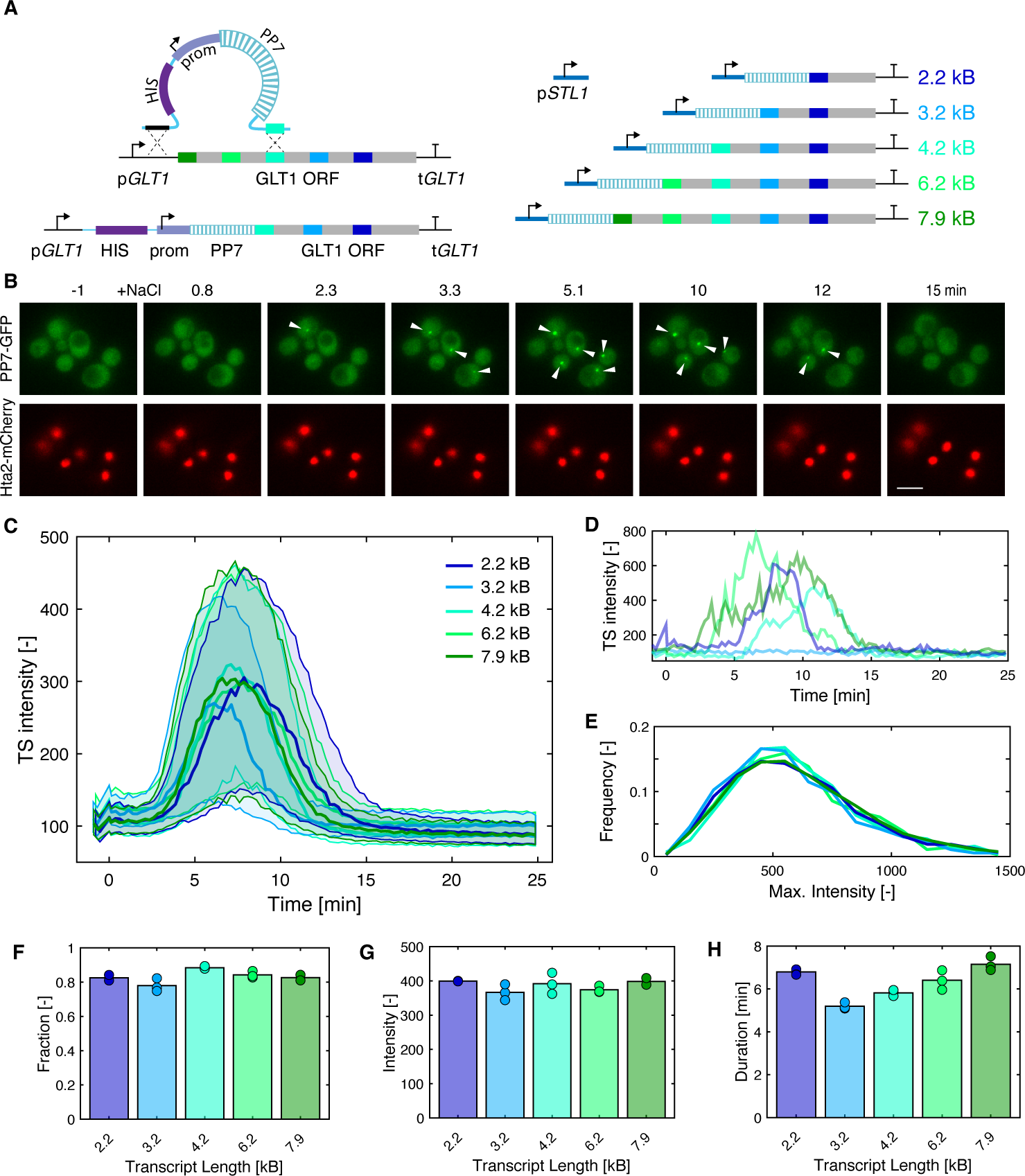
Effect of gene length on the quantification of nascent transcripts induced by the STL1 promoter. A. Scheme describing the strategy used to generate transcriptional reporters with similar genetic make-up using different homology recombination regions in the GLT1 ORF. B. Images of the PP7-GFP and Hta2-mCherry from a time-lapse movie. At time 0, the osmolarity of the medium increases to 0.2M NaCl. The transcription sites (marked with arrowheads) are generated by the induction of the p*STL1* promoter inserted upstream of the stem loop array. The scale bar represents 5µm. C. Median (solid line) and 25- and 75-percentiles (shaded area) of the population of cells stressed by 0.2M NaCl for constructs with lengths from 2.2 kB (dark blue) to 7.9 kB (dark green). D. Examples of four traces of transcribing cells with variable induction time or intensities and one non-transcribing cell (light blue). E. Histogram of the maximum intensity of the transcription site in the transcribing cells sub-population for transcript lengths from 2.2 kB (dark blue) to 7.9 kB (dark green). F. G. and H. Average of the fraction of responding cells (F), maximum intensity (G) and transcription site duration (H) as a function of transcript length. Each circle represents a biological replicate, and the bar represents the mean of the two to three replicates. Five hundred to a thousand individual cells were averaged for each replicate.

### Stress-dependent transcription

We generated an initial set of reporters using the p*STL1* hyper-osmotic stress responsive promoter, which is induced upon stimulation of the cells by addition of NaCl to the medium. Following this stress, an important upregulation of the transcription is observed for roughly 15 minutes (Figure 2B, C and D)^41,42^. For all five constructs, we observe a similar transcription pattern. The percentage of cells in which we detect a transcription site is around 80% (Figure 2F). Although there is a great variability in the level of transcription in individual cells (Figure 2D and E), on average, the maximum intensity of the transcription site trace does not seem to vary as a function of gene length (Figure 2E and G).

However, there is a trend suggesting that the duration of the transcription decreases with gene length from 7.5 min for 7.9 kB down to 5 min for the 3.2 kB construct (Figure 2H). The smallest 2.2 kB transcript is excluded from this trend. The formation of aggregates was observed with this construction, probably due to the small size of the truncated GLT1 mRNA relative to the GFP decorated PP7 stem loops (Figure S1). These fluorescent foci are freely moving in the cytoplasm and can be falsely interpreted as transcription sites in our automated analysis pipeline extending the measured duration of transcription. Taken together, the independence of the signal intensity from transcript length and the variation in duration of the signal suggest that p*STL1* induces a single burst of transcription as described in Figure 1D. To confirm this possibility, we needed to assess the time required by the polymerase to transcribe the whole locus.

### Elongation rate

The rate at which polymerases transcribe a gene has been evaluated by different methods and elongation speeds ranging from 10 to 100 bp/s have been measured^43–47^. Since this speed is controlled by numerous parameters, including promoter identity, genomic region, or cellular state^44^, we set out to measure this rate in our system using a combination of PP7 and MS2 loops spaced by 0, 2 or 4 kB^16,46^. The PP7 loops are produced first and bound by a PP7-mCherry construct, followed by MS2 loops that are synthesized and targeted by the MS2-GFP protein (Figure S2A and B). The speed at which the polymerase travels along the gene can be estimated from the difference between the appearance of the red and green signals. Using our automated image analysis pipeline, the cells with transcription sites in both red and green channels were selected (Figure S2C). Then, manual curation was used to precisely detect the first frames in the time-lapse where the fluorescent signal accumulated at the transcription site (Start Time) (Figure S2D). The difference in Start Time evaluated in more than 80 individual cells per strain was used to calculate an average elongation speed of 63 bp/s which is identical for the 2 kB or 4 kB spacers (Figure S2E).

Based on this elongation speed, transcription of a 7.9 kB transcript should take approximately 2 minutes versus 50 seconds for a 3.2 kB transcript. Experimentally, the duration of the PP7 signal lasts on average 8 and 5 minutes for the 7.9 and 3.2 kB constructs, respectively. Relative to the expected duration, these values suggest that the observed transcriptional pulse is not generated by a single transcriptional burst, but rather that multiple rounds of transcription contribute to the measured signal. These multiple overlapping transcription events should result in a larger number of active polymerases accumulating on the longer transcript relative to shorter ones. Surprisingly, our measurements performed with the p*STL1* promoter do not indicate an increase in intensity as a function of gene length.

### Other promoters

The relationship between gene length and transcription site intensity signal was further monitored for a short and long genes controlled by different promoters. Two additional stress-inducible promoters p*GPD1* and p*HSP12* were tested (Figure 3A and B, Figure S3A and B)^48^. While the dynamics and intensity of the transcription site signal are different for each promoter upon hyper-osmotic shock, we observe that the short and long transcripts generate very similar transcriptional dynamic signals.

**Figure 3.**
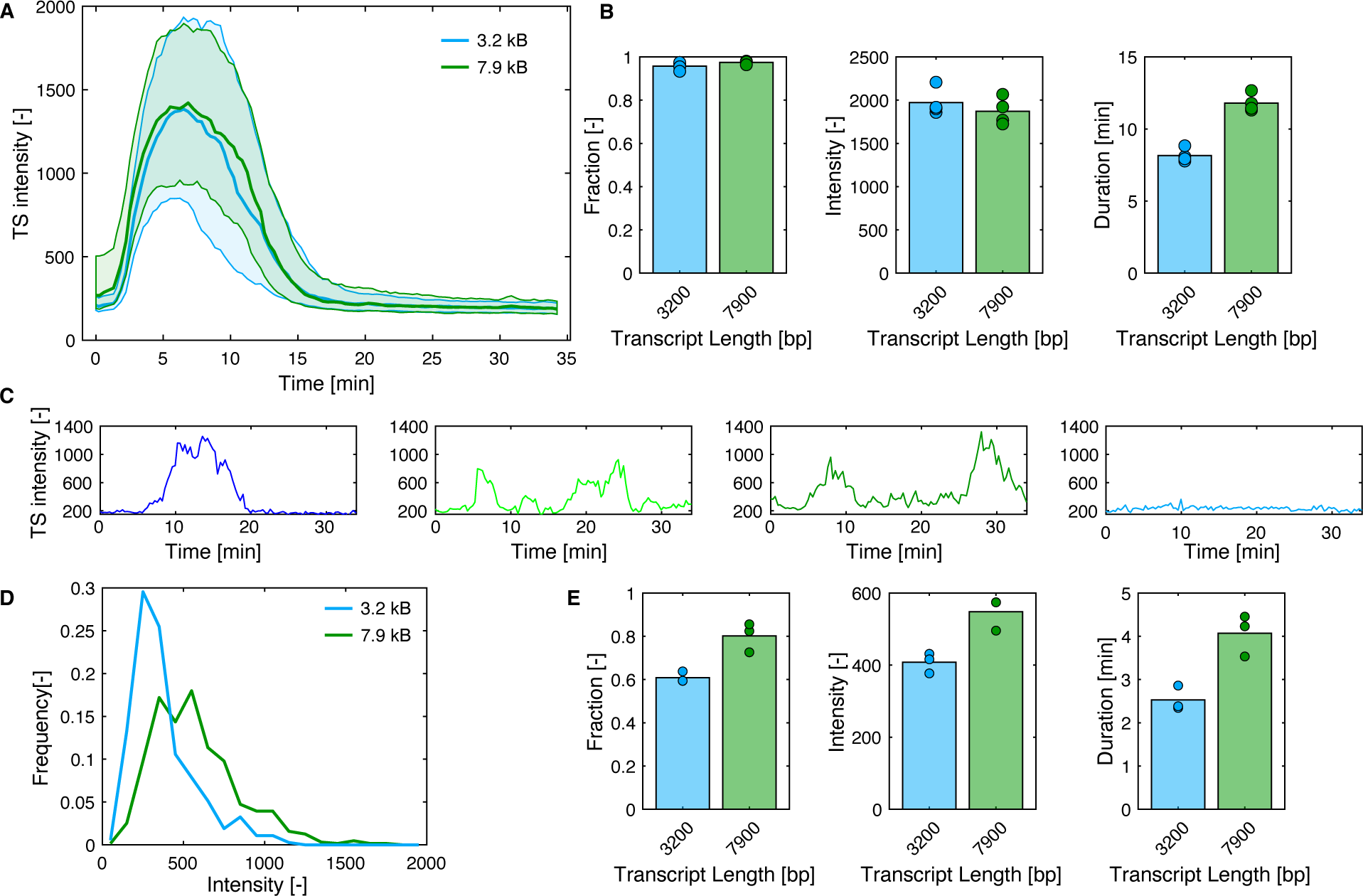
Induced and basal activity of the p*GPD1* promoter. A. Median (solid line) and 25- and 75-percentiles (shaded area) of the transcription site intensity of the population of the cells stressed by 0.2M NaCl for reporter constructs of 3.2 kB (blue) and 7.9 kB (dark green). B. Average of the fraction of responding cells, maximum intensity and transcription site duration as function of transcript length in induced conditions for the GPD1 promoter. Each circle represents a replicate, and the bar represents the mean of the three replicates. C. Single cell traces of the transcription site intensity observed for the basal activity of the GPD1 promoter. D. Histogram of the maximum intensity of the transcription site in the transcribing cells sub-population for transcript lengths for 3.2 kB (blue) and 7.9 kB (dark green). The maximum intensity is calculated from the difference between the three highest and three lowest data points in the trace. E. Average of the fraction of responding cells, maximum intensity and transcription site duration as function of transcript length for p*GPD1* in basal conditions. Each circle represents a biological replicate and the bar represents the mean of the three replicates.

In contrast to many other stress response genes which are strongly repressed under normal growth conditions, the GPD1 promoter possesses a substantial basal expression level. In vegetatively growing cells, p*GPD1* is activated stochastically and at relatively high levels in a large fraction of the population (Figure 3C and D). Interestingly, under these conditions, we observe a clear difference in transcription site intensity between short and long transcripts (Figure 3D and E). Moreover, the fraction of cells in which we detect a transcription site is lower for the short transcript (Figure 3E). Because the two promoters are inserted in the same genomic environment, their activity should be similar. The fact that we measure fewer transcriptionally active cells with lower intensities for the short transcript suggests that some of the transcriptional events fall below our detection threshold. In agreement with the scheme presented in Figure 1C, we believe that the signal from fewer nascent transcripts is accumulated on the short transcript, explaining the difference observed between the two reporters.

In parallel, we also quantified the transcriptional induction of the mating promoter p*AGA1* ^49,50^. In contrast with the stress-inducible constructs, this promoter displays a large difference in the transcription site intensity between the short and long transcripts (Figure S3C and D). The fraction of transcribing cells detected is lower for the short transcript. If we consider only the transcribing cells, the mean transcription site intensity remains 30% lower for the cells bearing the short versus the long reporter gene (Figure S3E). These data demonstrate the large difference in signal that can be observed between long and short transcripts.

Therefore, the relationship between transcription site intensity and transcript length seems to depend on the identity of the promoter. While the stress-induced promoters p*STL1* and p*HSP12* display a signal that is independent of gene length, the mating induced promoter p*AGA1* shows a strong dependence on gene length. Moreover, depending on conditions, the same promoter p*GPD1* will display an influence of gene length on transcriptional site intensity in vegetatively growing cells, while no dependence is observed under hyper-osmotic stress conditions. Taken together, these results suggest that biological parameters, which vary depending on the promoter identity and the cellular state, influence the relationship measured between gene length and transcription site intensity.

### Model of transcription

To understand which parameters could influence the relationship between gene length versus transcription site intensity, a basic model of osmotic stress response transcription was established (Figure S4). Osmotic stress and the ensuing adaptation driven by the HOG pathway were approximated by a step induction and an exponential decay. The initiation of transcription was controlled by the activation of a transcription factor (TF) by the stress. The active TF subsequently binds to the DNA. Next, the complex between the TF and DNA recruits the polymerase to the promoter. Once bound on the gene, the polymerase will start to transcribe the downstream sequence. In order to simulate the production of nascent mRNAs, the gene was split into 100 bp segments. A stochastic solver was used to simulate the progression of the polymerase on the ORF from the promoter to the terminator. Once the polymerase reaches the terminator, the mRNA is released, and the polymerase returns to the pool of free polymerases. For each polymerase advancing on the ORF, a fluorescent phage coat protein signal was calculated, recapitulating the expected transcription site intensity signal produced on the locus of interest by our reporter system. Some rate constants of the model (Table S5) are based on values that could be extracted from the literature^16,51,52^, while other parameters were adjusted to generate the expected number of mRNAs produced upon hyper-osmotic shock^53^.

Performing a thousand stochastic runs of this model enabled simulation of the diversity in transcriptional behavior between single cells. The means of the simulated transcription site intensities are displayed in Figure 4A along with the distribution of the maxima of the transcription site intensity and a histogram of the number of transcripts produced in each stochastic run. The simulations were performed with gene lengths varying from 2 to 8 kB (i.e. 20 to 80 polymerase steps along the locus). The first implementation of this model (elongation-dependent) based on these simple assumptions predicts a strong dependence between the transcription site signal and gene length as we initially anticipated (Figure 1C). However, these modeling results disagree with the experimental data obtained for the stress-inducible promoters (Figure 2C, 3A and S3A).

**Figure 4.**
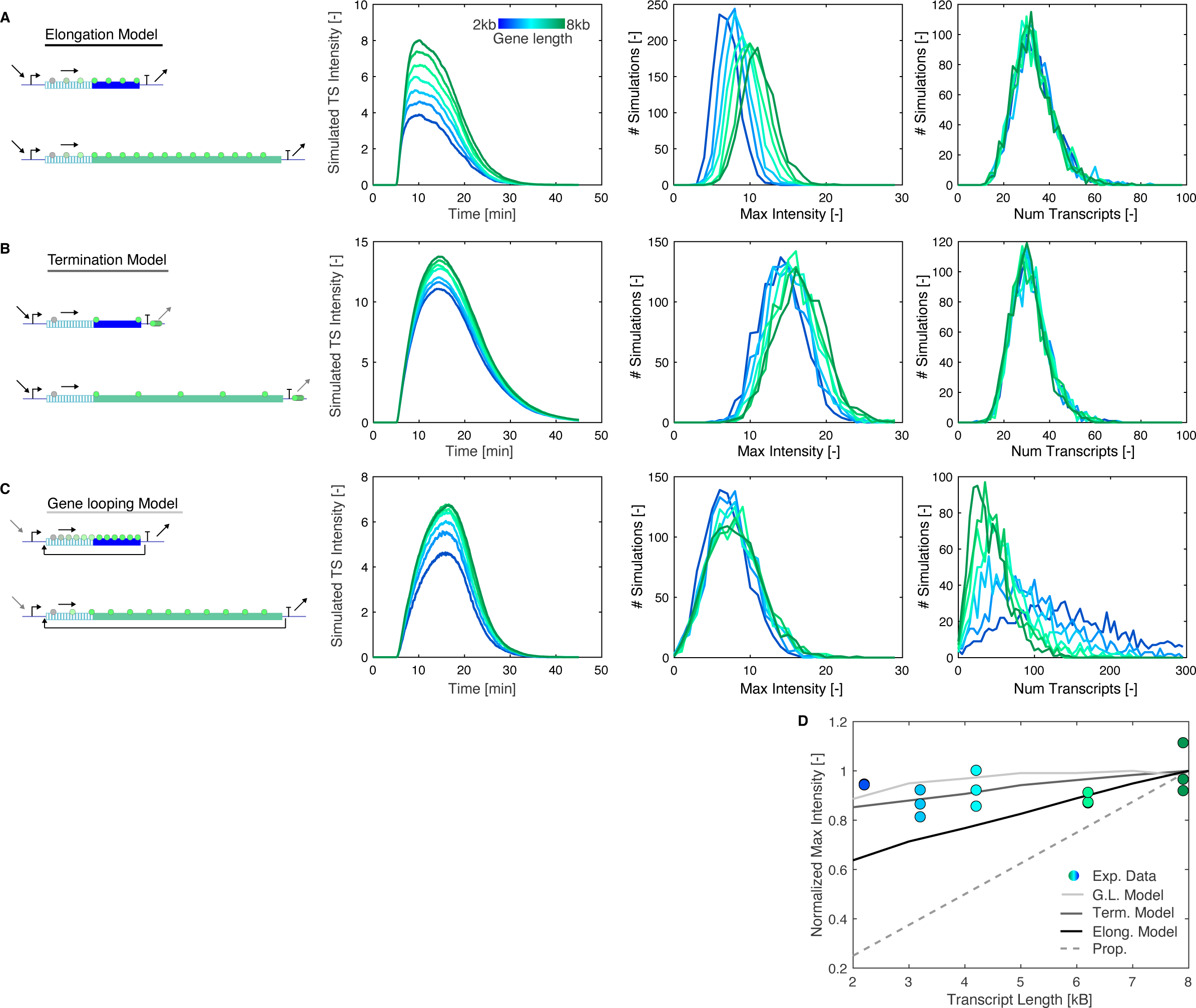
Results of the simulation of three model variants of stress-induced transcription. A. The elongation-dependent model predicts a strong dependence of the maximum signal intensity with gene length. In panels A, B and C, the left plot represents the mean simulated transcription site intensity from the 1000 stochastic runs. The central plot is a histogram of the maximal transcription site intensity reach in each simulation. The right plot is a histogram of the total number of mRNAs produce in each run of the simulation. B. The termination dependent model has a slow termination step (gray arrow) which leads to an accumulation of transcripts at the terminator and a lower dependence of the transcription site intensity with gene length. C. The gene looping model includes a recycling of the polymerase from the terminator to the promoter. It is also characterized by a lower activity of the promoter (gray arrow). D. Comparison between the predicted and measured variation of maximum intensity of the transcription site with gene length. The solid lines represent the simulation data. The dots are the normalized transcription site intensities measured for the p*STL1* induced transcription. The dashed line represents a linear relationship between transcription site intensity and gene length.

To reconcile the modeling output and the experimental findings, we set out to identify which parameters could influence the relationship between gene length and signal intensity. Different elongation speeds between short and long genes could in principle explain this behavior. It has been described that the polymerase initiate transcription at a slow rate and becomes more processive as it moves along the gene ^43^. In contrast, the accumulation of DNA supercoiling could slow down the transcription on longer genes^54^. To display a similar relative intensity on an 8 kB and a 3 kB gene, a polymerase on a 3 kB gene should have on average a 2.5 slower speed. Our elongation speed measurements with the 2 kB and 4 kB spacer rather suggest that the elongation speed is not dependent on gene length.

Polymerase pausing, if it happens homogeneously throughout the gene length, should also rather increase the signal for the long genes then decrease it because the residence time of the polymerase on the long genes should be increased. Note that with our 4 kB spacer we have observed two instances of pausing (out of 93 single cells quantified, Figure S2D) which lasted on the order of 4 minutes. All the other traces indicated a strong temporal correlation between the apparition of the RFP and the GFP foci. Early termination events could contribute to the observation of a relatively larger signal for short genes versus long ones. However, with the dual stem loop reporter, we observe only rarely (<3%) an RFP transcription focus without the presence of the subsequent GFP focus. Taken together, the measurements performed with PP7-MS2 coupled reporter suggest that once initiated, the transcription happens in a highly processive manner at a relatively constant speed with rare pauses and exceptional early termination events.

However, one parameter that could influence the relative intensity of short versus long genes is the rate of termination. If a nascent mRNA transcript lingers on a locus for an extended period of time due to a slow termination process, the time spent by the polymerase on genes of various lengths becomes comparatively less important. The results of the simulation for this termination-dependent model indeed show that the transcription site intensity signal becomes less dependent on gene length (Figure 4B).

An alternative explanation for the experimental behavior observed is the establishment of a gene loop^2,55–57^. Gene loops are formed by a close interaction between the promoter and terminator of a gene, via the association of the 3’-end processing machinery and the general transcription factor (TFIIB)^55^. Formation of the loop has been shown to enhance transcriptional memory and favor directionality of transcription^58,59^. One additional role of this complex formation is to recycle the polymerase from the terminator to the promoter to initiate a new transcription of the locus^2,60,61^. In order to simulate this phenomenon, the model was slightly modified to include an additional reaction. When a polymerase reaches the terminator, it can be recycled back to the start of the gene instead of being sent back to the pool of free polymerase (Figure S4). This transfer from the terminator to the promoter was modeled as a stress-dependent reaction, to prevent continuous transcription once stress adaptation has occurred.

In all model variants, the number of polymerases loaded on the gene by the promoter is independent of gene length. However, in the gene looping model variant, polymerase recycling occurs more frequently on shorter genes because polymerases reach the terminator more rapidly on short genes than on long ones, resulting in a higher density of polymerases acting on short genes. The nascent mRNA signal intensity is equivalent for short and long genes because the overall number of polymerases acting on the loci is similar. (Figure 4C). A direct implication of this mechanism is that the number of mRNA produced by a short gene is higher than for a longer transcript.

### Pulsatile activation

Two different variants of the model can provide a valid explanation for our experimental measurements: slow termination or formation of gene loops (Figure 4D). One possible way to distinguish these two variants is to test the response of the cells to a transient activation of the pathway. Indeed, one prediction from our model is that if the termination is a slow process, the transcription site will have a long half-life even if the stimulus is shutoff, unlike for the elongation and gene loop model variants (Figure 5A).

**Figure 5.**
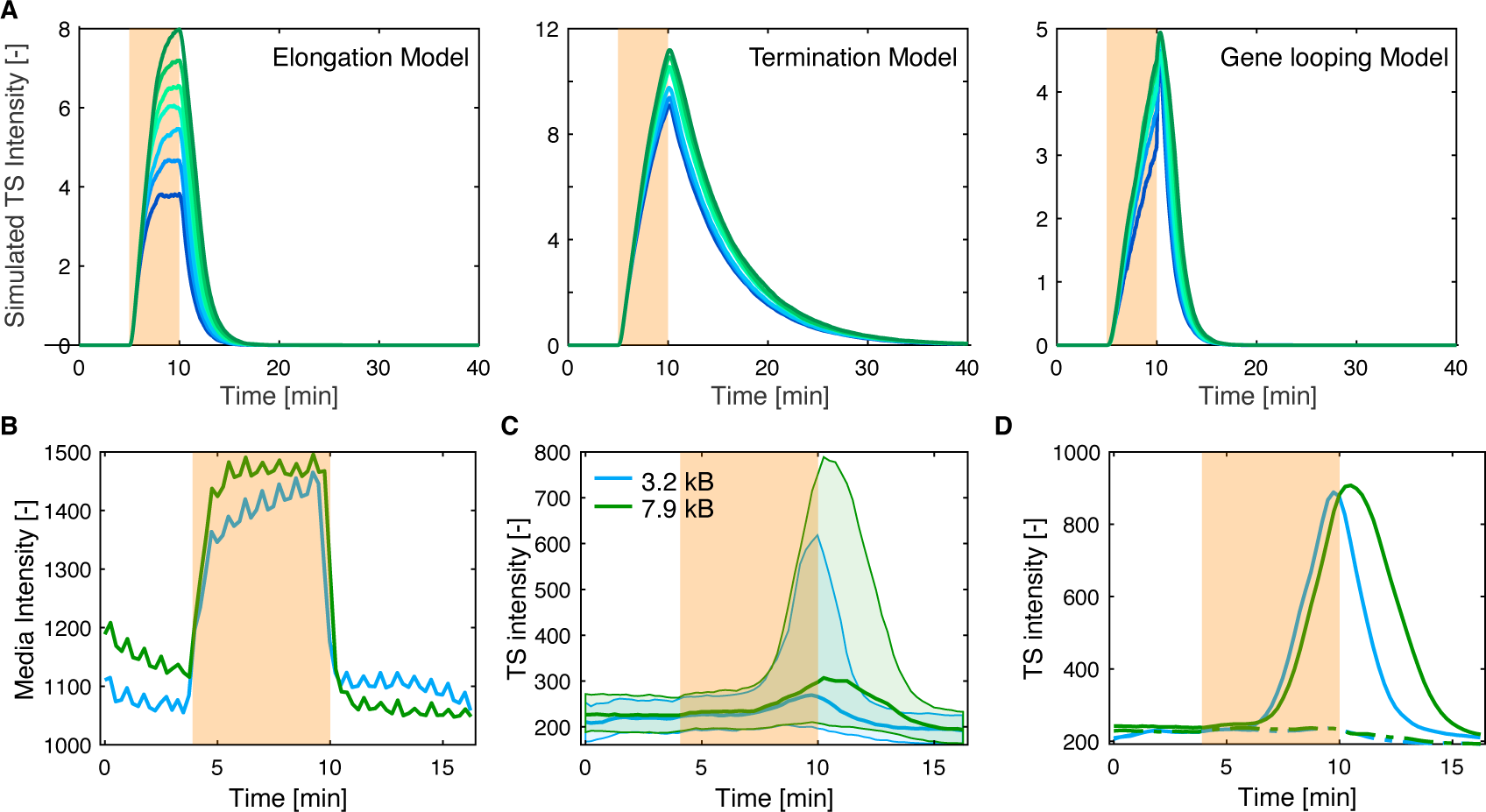
Pulsatile activation of the HOG pathway. A. Simulation of the transcription site intensity dynamics for the three model variants when a transient activation of the pathway is used (orange area). The Elongation and Gene Looping models predict a rapid decrease of the signal once the stimulus stops while the termination model displays a long-lasting signal. B. Pulse of 0.2M NaCl applied on the cell population in a flow chamber quantified by the level of fluorescence in the medium. C. Median (solid line) and 25- and 75-percentiles (shaded area) of the transcription site intensity of the population of the cells stressed by the transient 0.2M NaCl pulse for transcription constructs controlled by the p*STL1* promoter with lengths of 3.2 kB (blue) and 7.9 kB (dark green). D. Average transcription site intensity across the sub-population of responding (solid line) and non-responding cells (dashed line) for the 3.2 kB (blue) and 7.9 kB (dark green) constructs. The orange area in panels B, C and D highlight the time when the high osmolarity medium is present.

If a brief NaCl pulse stimulates the cells, Hog1 activity quickly returns to a basal level once the stress is removed and the induction of the STL1 promoter will stop^42,62^. According to our model, if termination is a slow process, the produced transcript can linger for up to 10 to 15 minutes at the transcription site. To test this hypothesis, we used flow chambers to stimulate the cells with a 7-minute pulse of 0.2M NaCl (Figure 5B). The transcription site intensity starts to rise 3 minutes after the beginning of the pulse. When the conditions are shifted back to the low osmolarity medium, the transcription site intensity starts to decline (Figure 5C and D). Five minutes later the transcription sites have disappeared. This behavior contradicts the existence of a slow termination process and instead demonstrates the highly dynamic nature of the transcription site visualized with the fluorescent PP7. These data also exclude the possibility that non-physiological aggregates of PP7 bound transcript are formed at the transcription site and preclude our observation of the true transcriptional dynamics.

The short 3.2 kB and long 7.9 kB genes display an important difference in the decay of the PP7 signal, which amounts to almost 1 and a half minute (Figure 5D). If one considers a polymerase that has initiated transcription just before the hyper-osmotic stress is relieved and subsequently completes the transcription of the locus, the signal from this nascent transcript is expected to remain present longer on the 7.9 kB gene than on the short one. Based on our measured elongation speed, a difference of 1 minute and 15 seconds between the two reporters is expected, which is in line with the pulse response measurements.

### Gene looping

Since the pulse experiments invalidated the termination-dependent model, we next wanted to verify the gene looping hypothesis. As shown in Figure 4, one specific feature of the gene looping-dependent model is that smaller genes produce more transcripts due to more prevalent recycling of the polymerase on these constructs. In the two other model variants, the number of transcripts produced is strictly dependent on the promoter activity and remains independent of gene length.

To test this prediction, we quantified by RT-qPCR the amount of mRNA produced from the GLT1 locus in strains where the gene was modified by insertion of the p*STL1*-PP7sl constructs at various positions. We compared these measurements to the number of transcripts expressed from the endogenous STL1 and CTT1 stress-response genes. We evaluated the amount of mRNA produced as a function of time in strains bearing the 3.2 and 7.9 kB reporters. The two endogenous stress-response genes are transiently induced upon 0.4M NaCl stimulus in YPD with a peak of production at 10 min. The measured response is largely comparable between the two strains for these two loci (Figure 6A). For the GLT1 locus, primers were chosen at the end of the GLT1 ORF which is common to both strains. The production from the short transcript is significantly higher at early time points. At 30 minutes, the difference is no longer significant, possibly due to an increased stability of the longer mRNA.

**Figure 6.**
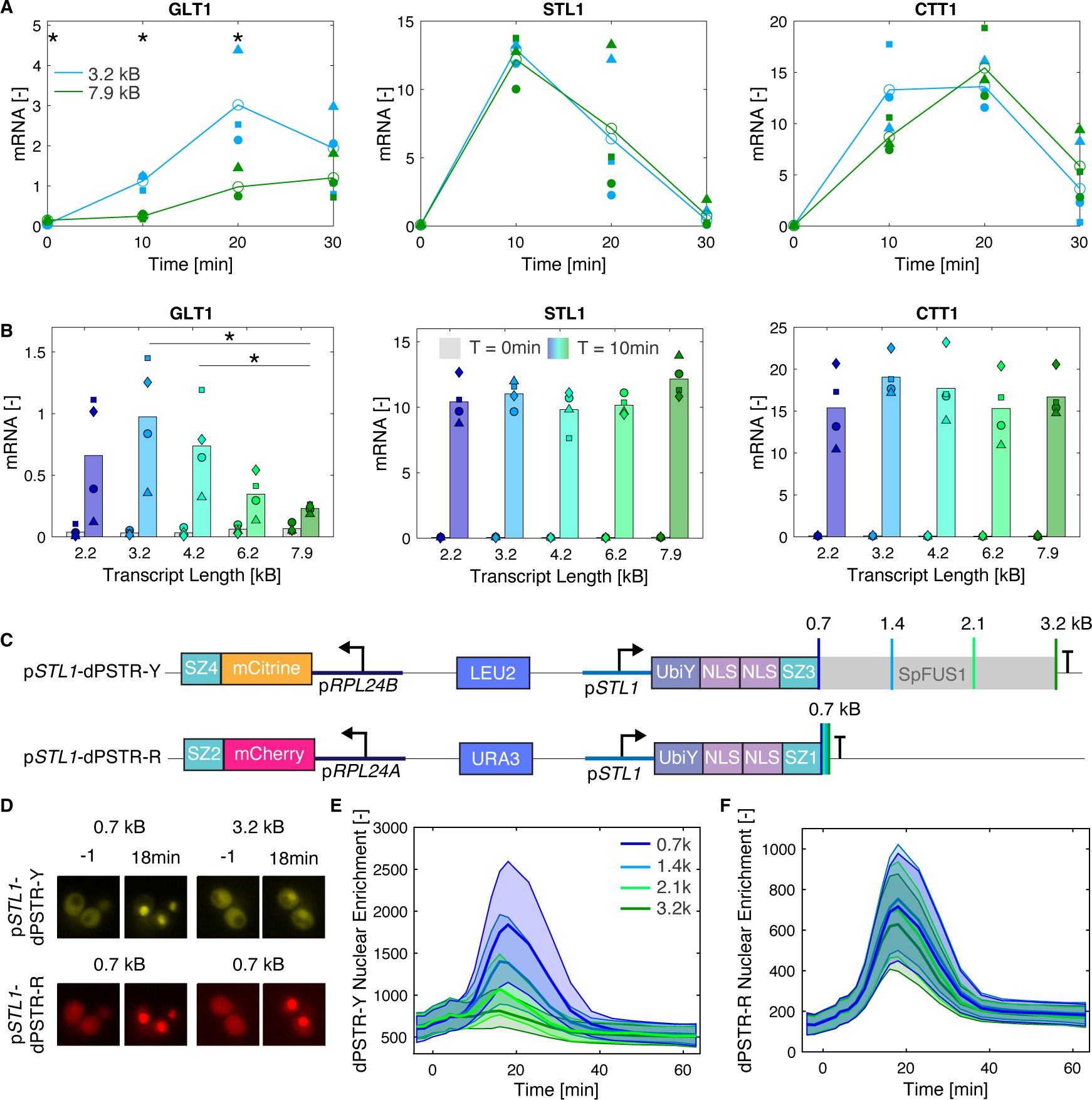
Influence of the gene length on mRNA and protein levels. A. Time course measurements of mRNA levels for the GLT1, STL1 and CTT1 ORF following 0.4M NaCl stress for the strains containing the 3.2 kB (blue) and the 7.9 kB (dark green) reporters. The open circles represent the averages of three biological replicates shown with the closed symbols. Symbols of the same shape designate the replicates from the same experiment. The asterisks indicate a significant difference between the two strains at the same time point. B. mRNA levels measured before (gray bars) and after stress (colored bars) for the GLT1, STL1 and CTT1 ORF following 0.4M NaCl stress. The bars represent the average of 4 biological replicate experiments and the symbols, the results from each replicate. Symbols of the same shape designate the replicates from the same experiment. The asterisks indicate a significant difference between the shorter transcripts relative to the 7.9 kB transcript. C. Scheme of the two transcriptional reports integrated in the LEU2 and URA3 loci. Each dPSTR consists of two transcriptional units: the inducible part and the constitutively expressed one, which includes the fluorescent protein. On the yellow dPSTR, the length of the inducible transcript has been extended by the insertion of *S. pombe* FUS1 sequences, but the ORF size remains the same. D. Thumbnail images of cells before and 18 minutes after 0.2M NaCl stimulation. The level of nuclear enrichment is a measure of protein expression of the inducible dPSTR moiety. E. and F. Quantification of the nuclear relocation of the fluorescent protein upon induction of the p*STL1*-dPSTR reporter in the yellow (E) and red channels (F). The relocation for the longer transcripts (green) is decreased relative to the shorter ones (blue), while the red expression reporter indicates a similar induction for all four strains.

A similar experiment was performed with the five strains with different lengths of GLT1 reporter constructs. The transcriptional induction was quantified 10 minutes after the stress. The amount of mRNA produced is clearly dependent on gene length (Figure 6B). These results are in line with the gene looping model which suggests that because of the more frequent recycling of the polymerase from the terminator to the promoter for short genes, an increased number of transcripts will be generated from the short genes.

The influence of gene length on protein production levels was also tested using a strain bearing two dynamic protein synthesis translocation reporters (dPSTR) in the yellow and red channels^33^. These reporters function using a constitutively expressed protein, whose enrichment in the nucleus is controlled by the production of a small inducible peptide. Both dPSTR sensors are under the control of the STL1 promoter; however, the yellow reporters encode the same inducible peptide on transcripts with length varying from 0.7 to 3.2 kB (Figure 6C). The induction of protein synthesis upon 0.2M NaCl stress is identical in all strains when quantifying the red signal. In comparison, the signal from the yellow reporter decreases fourfold when the transcript length increases from 0.7 to 3.2 kB (Figure 6D, E and F).

Taken together, these results show that the total amount of mRNA produced is significantly influenced by the gene length and thereby affects the quantity of protein produced. The RT-qPCR data indicate that more polymerases are active on the shorter genes and thereby validate our hypothesis that polymerase recycling is more prevalent on the short gene than on the long ones. This difference in mRNA production as a function of gene length is further relayed to the protein synthesis level, as observed with the dPSTR measurements. However, other parameters could also contribute to influence the final amount of proteins produced as function of transcript length such as the mRNA export, mRNA stability or translation efficiency.

To test how the propensity to form gene loops was encoded in a promoter, we fused the upstream activation sequence (UAS) of STL1 and the core promoter of p*AGA1* (Figure S5A). In this construct the regulation from the HOG pathway impinges on the UAS, but the general transcription factors which play a key role in the gene looping mechanism^60^ are recruited by the core region. In this context, we observe only a minor difference between the PP7 signal generated from a short (3.2 kB) or a long (7.9 kB) transcript (Figure S5B and C). These results are in line with the observation made with the GPD1 promoter which switches from a low fraction of gene looping to a higher one upon HOG pathway activation. These two experiments suggest that factors recruited via the UAS interact with the transcriptional machinery to enhance the formation of gene loops.

## Discussion

Intuitively, a dependence of the phage-coat protein signal at the transcription site on gene length is expected. This prediction applies well to the mating promoter p*AGA1*, where a large difference in the signal between the short and the long gene has been measured. This difference can be attributed to the longer residence time of the polymerase on the 7.9 kB gene, resulting in the integration of the fluorescent signal emanating from more nascent transcripts than on the short 2.2 kB gene. Astonishingly, this rule does not apply for the tested stress-inducible promoters that display an independence of transcription site intensity from gene length. Based on our mathematical simulations, two mechanisms could explain this unexpected relationship: slow termination or formation of gene loops. We dismissed the slow termination hypothesis because a transient activation of the pathway with a pulse of stress medium results in a fast disassembly of the transcription site foci. This is difficult to account for if termination is governed by a slow reaction rate.

The other hypothesis we can envision is the formation of gene loops. It has been demonstrated that gene loops are present on a large variety of genes in yeast^63^ and higher eukaryotes^64,65^. One function of this structure is to favor the recycling of polymerase from the terminator to the promoter^2^. From this scenario, we predict that the amount of mRNA produced by a short gene should be higher than by a long gene. We validated this assumption by using RT-qPCR to quantify the total amount of mRNA produced from a short or long gene in the same genomic environment. Interestingly, these data also highlight the difference between snapshot measurements, such as RT-qPCR or FISH, and phage coat protein assays. The latter provide a measure of the instantaneous polymerase activity on a locus, which is not straightforward to translate into a measure of total mRNA production. Temporal integration over the dynamic fluorescence signal can provide a relative measure of transcriptional output for reporters with the same genomic architecture. However, precise quantification of the elongation speed, termination rate and conversion of the fluorescent signal in a number of active polymerases^66^ would be required for a complete characterization of the transcriptional output.

For the elongation and termination model variants, each polymerase loaded on the gene via the promoter produces a single transcript. In the gene looping model variant, each polymerase can undergo multiple rounds of transcription. In order to produce mRNA in quantities matching experimental observations ^53^, we had to reduce the inducibility of the promoter to compensate for the recycling of polymerases. Therefore, the equilibrium constants between active and inactive TF and DNA-bound versus unbound TF are decreased by a factor of 400, considerably reducing the likelihood of recruiting an active polymerase to the gene. This behavior is in line with the known switch-like behavior of stress response genes that are repressed by chromatin under normal growth conditions and become highly transcribed once induced^41^. The establishment of a gene loop to promote polymerase recycling could thus contribute to the high inducibility of stress promoters. In addition, the formation of this complex would also allow cells to spare precious resources such as transcription factors, which are known to have low abundance ^67^. TFs could be involved only in the initiation of transcription; once the polymerase starts transcribing, the formation of gene loops could allow generation of numerous transcripts with limited involvement of TF bound to the upstream-activating sequence, and mostly requiring the general transcription factors associated to the core promoter ^61^.

When analyzing the transcription arising from the p*GPD1* promoter, we have observed that under inducing conditions there is no dependence of the signal on transcript length, while in the basal state the shorter gene produces a significantly lower signal compared to the long gene. These data suggest that the fraction of polymerase recycled by gene looping can be modulated as a function of the cellular state. Experiments performed with the promoter fusion, suggest that it is the factors recruited by the UAS that favor the gene looping mechanism, for instance, the MAPK Hog1 could stabilize the gene loop in order to enhance the number of transcripts produced from the locus. However, it could also be an indirect effect created by the chromatin environment that becomes more permissive in stress-induced conditions^41^ or an anchoring of the locus at the nuclear pores upon stress^58,68,69^ which thereby favor the formation of gene loops.

Given the strong impact that the formation of gene loops can have on the number of active polymerases on a gene and thus on the total amount of mRNA produced, the formation of this complex needs to be taken into consideration when measuring and modeling transcriptional processes. Most mathematical models of transcription focus on the loading of polymerases on the gene via the promoter. While this step is obviously essential, the transcriptional output might be determined to a larger extent by the recycling process. In our simple model of transcriptional stress response, for a 2 kB gene, 95% of the transcripts are generated due to the recycling of the polymerase. This percentage is probably lower *in vivo*. However, given the comparable level of the PP7 signal on the short and long genes in our experimental dataset, the contribution of polymerase recycling to the gene output is far from negligible.

Interpretation of the data generated by phage-coat protein assays also requires a careful assessment to detect the influence of polymerase recycling. A single continuous pulse of fluorescence signal could correspond to multiple successive rounds of transcription by polymerases recycled from the terminator to the promoter. Comparing the effective elongation speed to the duration of the pulse could provide a first hint at the presence of gene loops promoting polymerase recycling.

One major unanswered question is why the stress-responsive and mating promoters tested seem to have different propensities for forming gene loops or recycling polymerase. Both promoter types were selected from MAPK-induced genes that are upregulated upon extracellular stimuli. One putative explanation is that the recycling of polymerase might increase transcriptional noise, which is undesired in a cell-fate decision system such as the mating response^42,70^, but has little negative consequence on stress adaptation where output speed is prioritized. In any case, the mechanisms that regulate the level of recycling of the polymerases for different promoters or in different conditions remain to be investigated.

## Acknowledgements

We thank all members of the Pelet lab for input on the project, Mariona Nadal-Ribelles from the IRB in Barcelona for advice on the RT-qPCR protocol. The research has been funded by the Swiss National Science Foundation (Grant Nr: 31003A_182431) and the Department of Fundamental Microbiology and the University of Lausanne. We thank Mathieu Parron for technical assistance.

## Authors’ Contributions

OK and SP devised the study and performed the standard PP7 induction experiments. GL performed the elongation speed measurements. GL and BS performed the flow channel experiments. YD performed the RT-qPCR experiments. OK, YD and VV constructed plasmids and generated yeast strains for this study. SP performed the data analysis and wrote the paper.

## Data sharing and Data availability

The raw imaging data will be made available upon request. The quantified measurements will be made available on Zenodo (10.5281/zenodo.10441806).

## Supplementary Figure Legends

**Figure S1** Visualization of PP7 aggregates.

Cells bearing the p*STL1*-PP7sl-2kB reporter and expressing the PP7-GFPenvy construct were stimulated with 0.2M NaCl. Following the burst in transcription, aggregates start to form in the cytoplasm of a few cells (highlighted with the arrowheads). The scale bar represents 5µm.

**Figure S2** Quantification of the elongation speed.

A. Schematic of the transcriptional reporter used to quantify the elongation speed based on a tandem of PP7 and MS2 stem loops spaced by the *S. pombe* FUS1 sequence. As the polymerase travels on the gene, the nascent mRNA will be first labeled in red by the PP7-mCherry and subsequently in green by the MS2-GFP.

B. Images of transcription sites evolving from red to yellow (arrow heads) upon activation of the STL1 promoter by a 0.2M NaCl stimulus in diploid cells expressing the PP7-mCherry and the MS2-GFPenvy and containing the 2 kB spacer reporter.

C. Percentage of cells where a transcription focus was detected in the green and red channels.

D. Correlation between the transcription Start Times measured in red and in green in the same cell for the construct without spacer (dark blue), with the 2 kB spacer (light blue) and with the 4 kB spacer (green). The X=Y diagonal is shown with the line. More than 80 individual cells were quantified for each sample. The two red circles highlight cells where a long pause in the transcription has been observed.

E. Table displaying the mean difference in Start Times between the green and the red channels. The elongation speed is calculated from the size of the spacer divided by the difference in start times between the 2 or 4 kB constructs relative to the 0 kB one.

**Figure S3** Transcriptional activity measured from the HSP12 and AGA1 promoters.

A. Median (solid line) and 25- and 75-percentiles (shaded area) of the transcription site intensity arising from the HSP12 promoter of the population of the cells stressed by 0.2M NaCl for reporter constructs of 2.2 kB (dark blue) and 7.9 kB (dark green).

B. Average of the fraction of responding cells and maximum transcription site intensity as a function of transcript length for p*HSP12* in induced conditions. Each circle represents a replicate and the bar represents the mean of the three replicates.

C. Median (solid line) and 25- and 75-percentiles (shaded area) of the transcription site intensity arising from the AGA1 promoter of the population of the cells stimulated with 1µM α-factor for reporter constructs of 2.2 kB (dark blue) and 7.9 kB (dark green).

D. Average of the fraction of responding cells and maximum transcription site intensity as a function of transcript length for p*AGA1* in induced conditions. Each circle represents a replicate and the bar represents the mean of the three replicates.

E. Average Transcription Site intensity across the sub population of responding (solid line) and non-responding cells (dashed line) for 2.2 kB (dark blue) and 7.9 kB (dark green).

**Figure S4.** Scheme depicting the mathematical model of transcription.

Stress starts at 0 and is shifted to 100 after 5 minutes. It is subsequently degraded to represent the cellular adaptation. Red dashed lines represent the catalytic activation of specific reactions by the Stress. The transcription factor (TF) is activated by the stress and can thus bind the DNA where this complex recruits the polymerase. The polymerase will then advance on the gene which is decomposed in multiple elements until it reaches the terminator. Then, the mRNA is released and the polymerase can return to the free cellular pool. In purple, the gene looping reactions are described. Upon reaching the terminator, the polymerase can release its mRNA and be recycled at the beginning of the gene.

**Figure S5.** Transcriptional activity measured a fusion between the STL1-AGA1 promoters.

A. Scheme of the fusion promoter between the UAS of STL1 (-800 to -163) and the core promoter of AGA1 (-150 to 0).

B. Median (solid line) and 25- and 75-percentiles (shaded area) of the transcription site intensity arising from the fusion promoter of the population of the cells stimulated with 0.2M NaCl for reporter constructs of 3.2 kB (light blue) and 7.9 kB (dark green). The dashed green line represents the median of the endogenous STL1 promoter with 7.9 kB downstream transcript. Note that the imaging conditions are not identical then for the data presented in Figure 2.

C. Average of the fraction of responding cells maximum transcription site intensity and duration as a function of transcript length for the fusion promoter of pSTL1 in induced conditions. Each circle represents a replicate, and the bar represents the mean of the three replicates.

