## Supplementary material for "Implication of polymerase recycling for nascent transcript quantification by live cell imaging": Figure S1 to Figure S5 and Tables S1 to Table S5

Tables S1 to Table S5

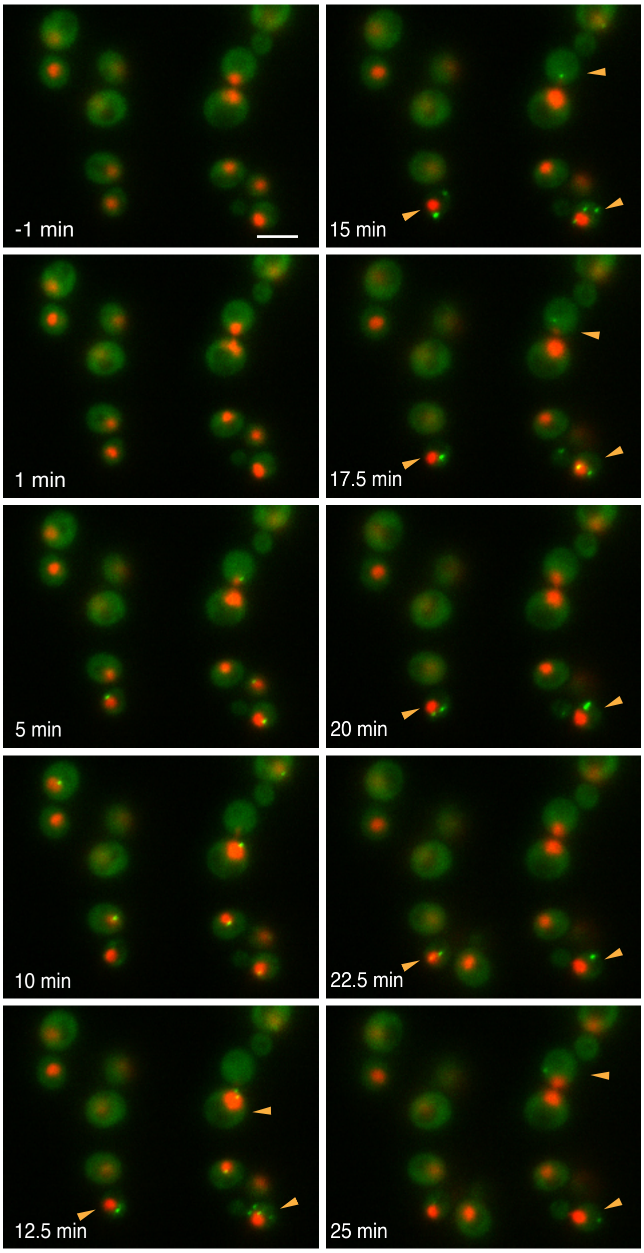

**Figure S1** Visualization of PP7 aggregates.

Cells bearing the p*STL1*-PP7sl-2kB reporter and expressing the PP7-GFPenvy construct were stimulated with 0.2M NaCl. Following the burst in transcription, aggregates start to form in the cytoplasm of a few cells (highlighted with the arrowheads). The scale bar represents 5µm.

A

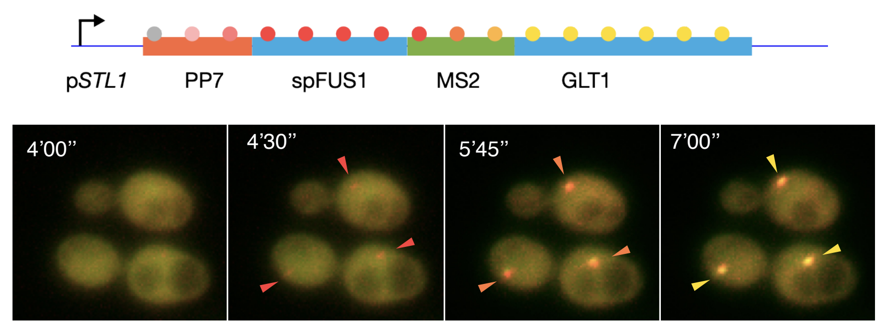

C

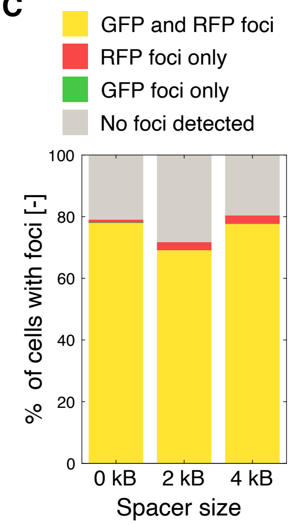

D

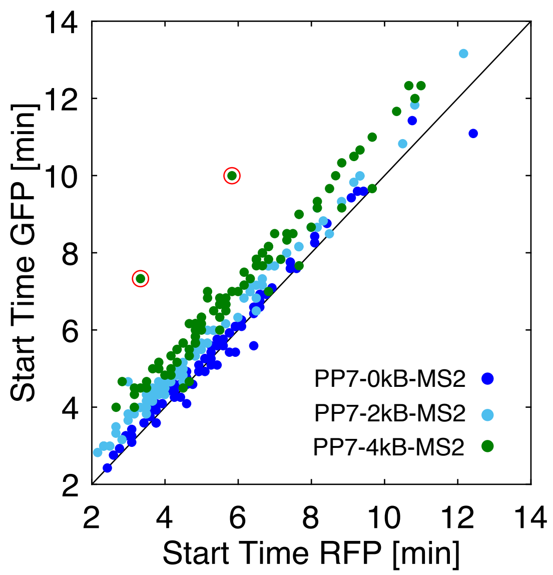

E

| | $\Delta T_{GFP\text{-}RFP}$ [s] | Elong. speed [bp/s] |
|---|---|---|
| PP7-0kB-MS2 | 5.8 | |
| PP7-2kB-MS2 | 37.6 | 63 |
| PP7-4kB-MS2 | 69.3 | 63 |

**Figure S2** Quantification of the elongation speed.

A. Schematic of the transcriptional reporter used to quantify the elongation speed based on a tandem of PP7 and MS2 stem loops spaced by the *S. pombe* FUS1 sequence. As the polymerase travels on the gene, the nascent mRNA will be first labeled in red by the PP7-mCherry and subsequently in green by the MS2-GFP.

B. Images of transcription sites evolving from red to yellow (arrow heads) upon activation of the STL1 promoter by a 0.2M NaCl stimulus in diploid cells expressing the PP7-mCherry and the MS2-GFPenvy and containing the 2 kB spacer reporter.

C. Percentage of cells where a transcription focus was detected in the green and red channels.

D. Correlation between the transcription Start Times measured in red and in green in the same cell for the construct without spacer (dark blue), with the 2 kB spacer (light blue) and with the 4 kB spacer (green). The X=Y diagonal is shown with the black line. More than 80 individual cells were quantified for each sample.The two red circles highlight cells where a long pause in the transcription has been observed.

E. Table displaying the mean difference in Start Times between the green and the red channels. The elongation speed is calculated from the size of the spacer divided by the difference in start times between the 2 or 4 kB constructs relative to the 0 kB one.

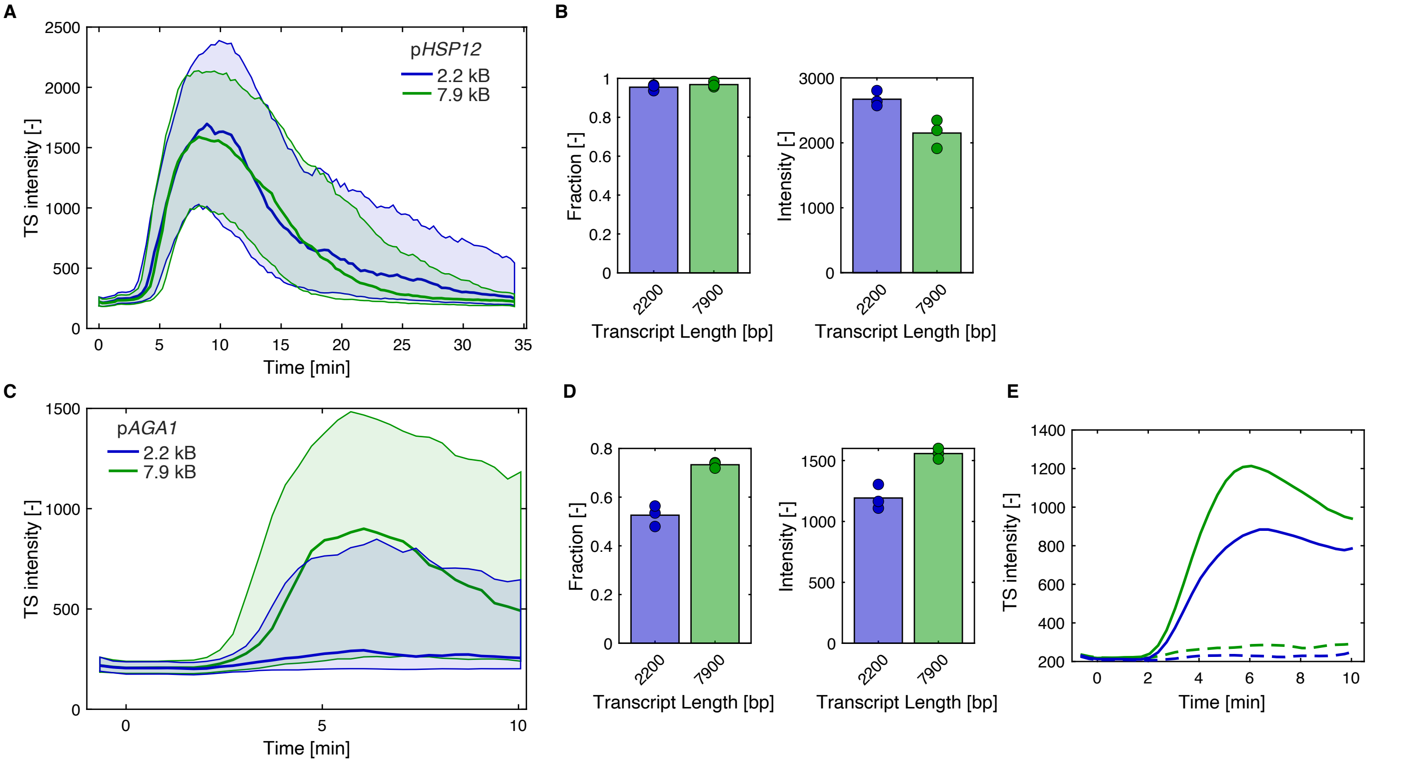

**Figure S3** Transcriptional activity measured from the HSP12 and AGA1 promoters.

A. Median (solid line) and 25- and 75-percentiles (shaded area) of the transcription site intensity arising from the HSP12 promoter of the population of the cells stressed by 0.2M NaCl for reporter constructs of 2.2 kB (dark blue) and 7.9 kB (dark green).

B. Average of the fraction of responding cells and maximum transcription site intensity as a function of transcript length for p*HSP12* in induced conditions. Each circle represents a replicate and the bar represents the mean of the three replicates.

C. Median (solid line) and 25- and 75-percentiles (shaded area) of the transcription site intensity arising from the AGA1 promoter of the population of the cells stimulated with 1µM α-factor for reporter constructs of 2.2 kB (dark blue) and 7.9 kB (dark green).

D. Average of the fraction of responding cells and maximum transcription site intensity as a function of transcript length p*AGA1* in induced conditions. Each circle represents a replicate and the bar represents the mean of the three replicates.

E. Average Transcription site intensity across the sub population of responding (solid line) and non-responding cells (dashed line) for 2.2 kB (dark blue) and 7.9 kB (dark green).

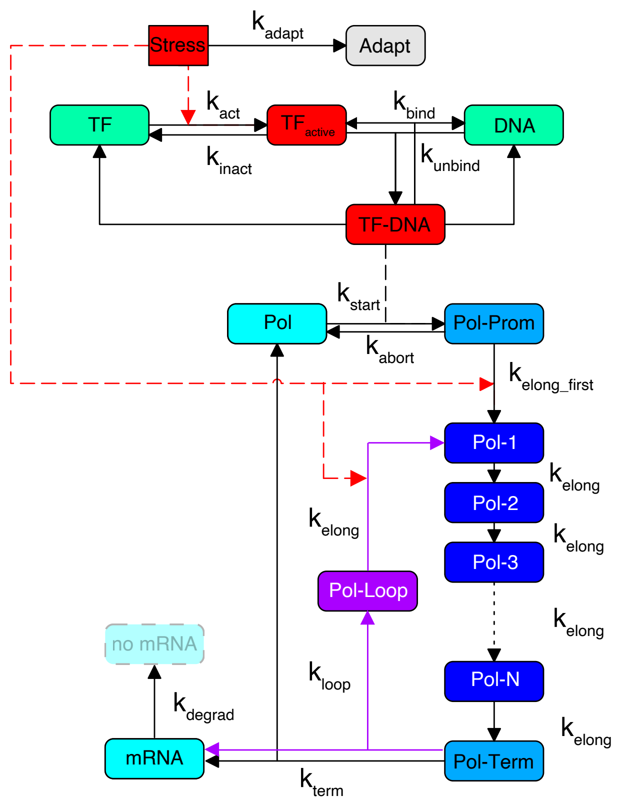

**Figure S4.** Scheme depicting the mathematical model of transcription.

Stress starts at 0 and is shifted to 100 after 5 minutes. It is subsequently degraded to represent the cellular adaptation. Red dashed lines represent the catalytic activation of specific reactions by the Stress. The transcription factor (TF) is activated by the stress and can thus bind the DNA where this complex recruits the polymerase. The polymerase will then advance on the gene which is decomposed in multiple elements until it reaches the terminator. Then, the mRNA is released and the polymerase can return to the free cellular pool. In purple, the gene looping reactions are described. Upon reaching the terminator, the polymerase can release its mRNA and be recycled at the beginning of the gene.

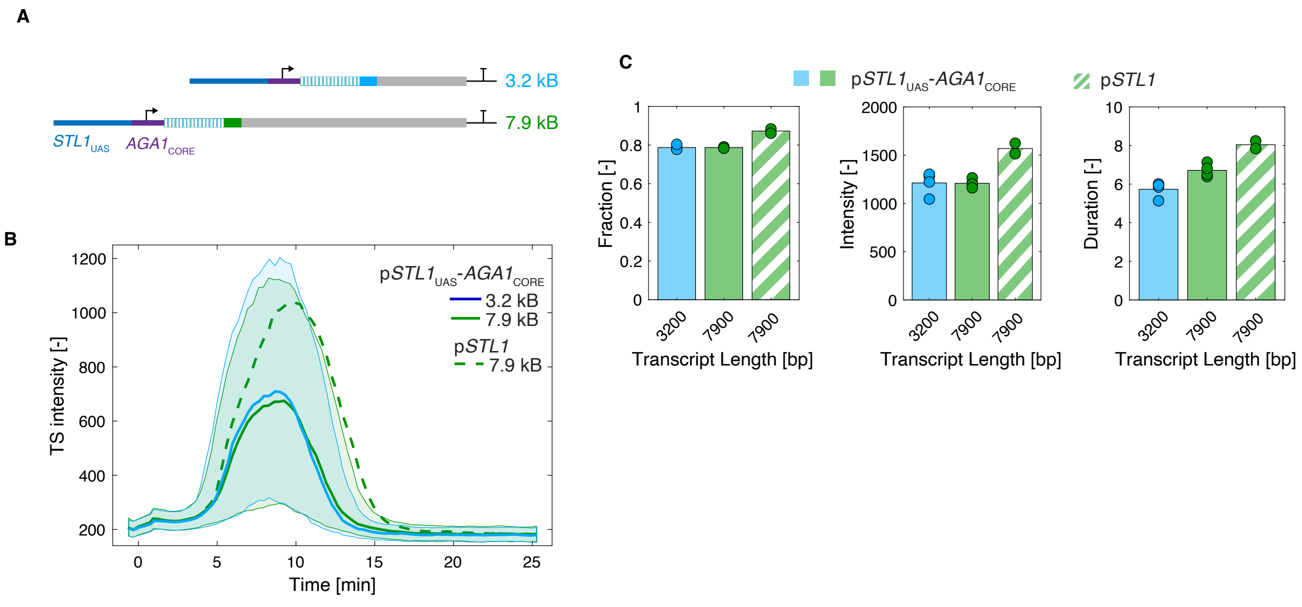

**Figure S5** Transcriptional activity measured a fusion between the STL1 and AGA1 promoters.

A. Scheme of the fusion promoter between the UAS of STL1 (-800 to -163) and the core promoter of AGA1 (-150 to 0).

B. Median (solid line) and 25- and 75-percentiles (shaded area) of the transcription site intensity arising from the fusion promoter of the population of the cells stimulated with 0.2M NaCl for reporter constructs of 3.2 kB (light blue) and 7.9 kB (dark green). The dashed green line represents the median of the endogenous STL1 promoter with 7.9 kB downstream transcript. Note that the imaging conditions are not identical then for the data presented in Figure 2.

C. Average of the fraction of responding cells maximum transcription site intensity and duration as a function of transcript length for the fusion promoter of pSTL1 in induced conditions. Each circle represents a replicate, and the bar represents the mean of the three replicates.

**Table S1. List of strains used in this study**

| Strain Name | Genotype | Plasmids | Ref. |
|---|---|---|---|
| ySP2 | W303 MAT**a** *leu2-3,112 trp1-1 can1-100 ura3-1 ade2-1 his3-11,15* | | 1 |
| yVW401 | ySP2<br>HTA2-mCherry:LEU2<br>pSIVu-pADH1-PP7-GFPenvy | pVW284 | 2 |
| yVW403 | yVW401<br>pHIS-pSTL1-PP7sl-GLT1 [7.9kB] | pSP264 | 2 |
| yOK1 | yVW401<br>pHIS-pSTL1-PP7sl-GLT1 [6.2kB] | pOK2 | |
| yOK2 | yVW401<br>pHIS-pSTL1-PP7sl-GLT1 [5.2kB] | pOK3 | |
| yOK3 | yVW401<br>pHIS-pSTL1-PP7sl-GLT1 [3.2kB] | pOK4 | |
| yOK4 | yVW401<br>pHIS-pSTL1-PP7sl-GLT1 [2.2kB] | pOK5 | |
| yVW432 | yVW401<br>pHIS-pGPD1-PP7sl-GLT1 [7.9kB] | pVW200 | 2 |
| yOK19 | yVW401<br>pHIS-pGPD1-PP7sl-GLT1 [3.2kB] | pOK19 | |
| yVW429 | yVW401<br>pHIS-pHSP12-PP7sl-GLT1 [7.9kB] | pVW293 | 2 |
| yOK20 | yVW401<br>pHIS-pHSP12-PP7sl-GLT1 [2.2kB] | pOK24 | |
| ySP1088 | yVW401<br>pHIS-pAGA1-PP7sl-GLT1 [7.9kB] | pSP409 | |
| yOK10 | yVW401<br>pHIS-pAGA1-PP7sl-GLT1 [2.2kB] | pOK21 | |
| ySP1038 | MAT**a**/MATα<br>HTA2-tdiRFP:TRP1<br>HTA2-tdiRFP:NAT<br>pSIVu-pADH1-PP7-mCherry<br>pSIVu-pADH1-MS2-GFPenvy<br>pHIS-pSTL1-PP7sl-MS2sl-GLT1 | pVW296<br>pSP561<br>pSP619 | |
| yGL52 | W303 MAT**a**/MATα<br>HTA2-tdiRFP:TRP1<br>HTA2-tdiRFP:NAT<br>pSIVu-pADH1-PP7-mCherry<br>pSIVu-pADH1-MS2-GFPenvy<br>pHIS-pSTL1-PP7sl-2k-MS2sl-GLT1 | pVW296<br>pSP561<br>pSP667 | |

| | | |
|---|---|---|
| yGL53 | W303 MAT**a**/MAT**α**<br>HTA2-tdiRFP:TRP1<br>HTA2-tdiRFP:NAT<br>pSIVu-pADH1-PP7-mCherry<br>pSIVu-pADH1-MS2-GFPenvy<br>pHIS-pSTL1-PP7sl-4k-MS2sl-GLT1 | <br><br><br>pVW296<br>pSP561<br>pSP668 |
| ySP761 | W303 MAT**a**<br>HTA2-CFP:HIS3<br>pSIVu pSTL1-dPSTR-R<br>pSIVl pSTL1-dPSTR-Y [0.7kB] | <br><br>pDA183<br>pDA199 |
| ySP1070 | W303 MAT**a**<br>HTA2-CFP:HIS3<br>pSIVu pSTL1-dPSTR-R<br>pSIVl pSTL1-dPSTR-Y [1.4kB] | <br><br>pDA183<br>pSP688 |
| ySP1071 | W303 MAT**a**<br>HTA2-CFP:HIS3<br>pSIVu pSTL1-dPSTR-R<br>pSIVl pSTL1-dPSTR-Y [2.1kB] | <br><br>pDA183<br>pSP689 |
| yOK24 | W303 MAT**a**<br>HTA2-CFP:HIS3<br>pSIVu pSTL1-dPSTR-R<br>pSIVl pSTL1-dPSTR-Y [3.2kB] | <br><br>pDA183<br>pOK27 |
| ySP1091 | yVW401<br>pHIS-p[STL1-AGA1]-PP7sl-GLT1 [7.9kB] | <br>pSP710 |
| ySP1092 | yVW401<br>pHIS- p[STL1-AGA1]-PP7sl-GLT1 [3.2kB] | <br>pSP711 |

1. Ralser, Open Bio. 2012
2. Wosika Nat. Com. 2020

**Table S2. List of plasmids used in this study**

| Plasmid # | Backbone | Insert | Ref |
|---|---|---|---|
| pVW284 | pSIVura | pADH1-PP7-ΔFG-GFPenvy | 1,2 |
| pSP264 | pHIS | pSTL1-24xPP7sl-GLT1{1:305} | 1 |
| pOK2 | pHIS | pSTL1-24xPP7sl-GLT1{1720:2220} | |
| pOK3 | pHIS | pSTL1-24xPP7sl-GLT1{3720:4220} | |
| pOK4 | pHIS | pSTL1-24xPP7sl-GLT1{4740:5240} | |
| pOK5 | pHIS | pSTL1-24xPP7sl-GLT1{5740:6240} | |
| pVW200 | pHIS | pGPD1-24xPP7sl-GLT1{1:305} | 1 |
| pOK19 | pHIS | pGPD1-24xPP7sl-GLT1{4740:5240} | |
| pVW293 | pHIS | pHSP12-24xPP7sl-GLT1{1:305} | 1 |
| pOK24 | pHIS | pHSP12-24xPP7sl- GLT1{5740:6240} | |
| pSP409 | pHIS | pAGA1-24xPP7sl-GLT1{1:305} | |
| pOK21 | pHIS | pAGA1-24xPP7sl- GLT1{5740:6240} | |

| | | | |
|---|---|---|---|
| pVW296 | pSIVura | pADH1-PP7-ΔFG-mCherry | 1 |
| pSP561 | pSIVura | pADH1-MS2-GFPenvy | 1 |
| pSP619 | pHIS | pSTL1-24xPP7sl-24xMS2-GLT1{1:305} | |
| pSP667 | pHIS | pSTL1-24xPP7sl-SpFUS1[2kB]-24xMS2-GLT1{1:305} | |
| pSP668 | pHIS | pSTL1-24xPP7sl-SpFUS1[4kB]-24xMS2-GLT1{1:305} | |
| pDA183 | pSIVura | p*STL1*-UbiY -2xSv40NLS- SynZip1-t*CYC1*<br>p*RPL24A*-mCherry-SynZip2-t*SIF2* | 3 |
| pDA199 | pSIVleu | p*STL1*-UbiY -2xSv40NLS- SynZip3-t*CYC1*<br>p*RPL24B*-MCitrine- SynZip4-t*NUP53* | 3 |
| pSP688 | pSIVleu | p*STL1*-UbiY -2xSv40NLS- SynZip3-SpFUS1[0.7kB]-t*CYC1*<br>p*RPL24B*-MCitrine- SynZip4-t*NUP53* | |
| pSP689 | pSIVleu | p*STL1*-UbiY -2xSv40NLS- SynZip3-SpFUS1[1.4kB]-t*CYC1*<br>p*RPL24B*-MCitrine- SynZip4-t*NUP53* | |
| pOK27 | pSIVleu | p*STL1*-UbiY -2xSv40NLS- SynZip3-SpFUS1[2.5kB]-t*CYC1*<br>p*RPL24B*-MCitrine- SynZip4-t*NUP53* | |
| pSP710 | pHIS | pSTL1$_{\{-800 - -163\}}$-pAGA1$_{\{-150 - 0\}}$-24xPP7sl-GLT1{1:305} | |
| pSP711 | pHIS | pSTL1$_{\{-800 - -163\}}$-pAGA1$_{\{-150 - 0\}}$-24xPP7sl-GLT1{4740:5240} | |

1. Wosika Nat. Com. 2020
2. Wosika Mol. Gen. Gen. 2016
3. Aymoz Nat. Com. 2016

**Table S3. List of primers to amplify the GLT1 fragments**

| Gene | Forward | Reverse | Ref |
|---|---|---|---|
| GLT1 {1720:2220} | GTATCGGCTAGCCTCCAATTGACCCAATTCG | GTATCGGACGTCCCATGGCTAAGTATGGATAAACA | |
| GLT1 {3720:4220} | GTATCGGCTAGCCGAAATGGTGGGTCATTCT | GTATCGGACGTCCCATAGAAACAAGTGTTACCAACA | |
| GLT1 {4740:5240} | GTATCGGCTAGCCCAAGAAGTTGATGACGAAG | GTATCGGACGTCGTGTACAAGCTCCCTCAC | |
| GLT1 {5740:6240} | GTATCGGCTAGCGGAAATCATTCGTGAAAAGATCC | GTATCGGACGTCGAGTATGAGGAGTCGTCG | |

**Table S4. List of primers for the qPCR**

| Gene | Forward | Reverse | Ref |
|---|---|---|---|
| VCX1 | TGCGTGTGCATCCCTACTGA | AAGTGGTCTTCCTTGCCATGA | 1 |
| CTT1 | CATACGCCGCTCCATACCAGAA | CGAACAAGACCAGGACATTTGT | 1 |
| STL1 | AAGCATACGAGGATGGCACT | TCGTCGTCATAAGAGCCCAA | |
| GLT1 | TCAATGGCAACGATAACGAA | ACGTAGTGCCGTCCATTAGG | |

1. Klopf, NAR, 2016

**Table S5. Parameters used in the mathematical model**

| Constant | Value | Ref |
|---|---|---|
| $k_{adatp}$ | 1/3/60 | |
| $k_{act}$ | 0.05/Slow | |
| $k_{inact}$ | 0.01*Slow | |
| $k_{bind}$ | 0.05/Slow | |
| $k_{unbind}$ | 0.01*Slow | |
| $k_{start}$ | 0.01/PolConc | |
| $k_{abort}$ | 0.05*Slow | |
| $k_{elongfirst}$ | ElongSpeed/SegmentSize | |
| $k_{elong}$ | ElongSpeed/SegmentSize/StressMax | |
| $k_{loop}$ | ElongSpeed/SegmentSize | |
| $k_{term}$ | 0.015 or 0.005 | 1 |
| $k_{degrad}$ | 0.005 | 2 |
| ElongSpeed | 60 bp/s | |
| Slow | 0.01 or 2 | |
| PolConc | 100 | |
| StressMax | 100 | |
| Stress | 0 | |
| TF | 1 | |
| DNA | 1 | |

1. Larson Science 2011
2. Neuert Science 2013, Chan eLife 2018
